# Sialic acid and fucose residues on the SARS-CoV-2 receptor binding domain modulate IgG reactivity

**DOI:** 10.1101/2022.01.20.477056

**Authors:** Ebba Samuelsson, Ekaterina Mirgorodskaya, Kristina Nyström, Malin Bäckström, Jan-Åke Liljeqvist, Rickard Nordén

## Abstract

The receptor binding domain (RBD) of the SARS-CoV-2 spike protein is a conserved domain and a target for neutralizing antibodies. We defined the carbohydrate content of recombinant RBD produced in different mammalian cells. We found a higher degree of complex type N-linked glycans, with less sialylation and more fucosylation, when the RBD was produced in Human embryonic kidney cells compared to the same protein produced in Chinese hamster ovary cells. The carbohydrates on the RBD proteins were enzymatically modulated and the effect on antibody reactivity was evaluated with serum samples from SARS-CoV-2 positive patients. Removal of all carbohydrates diminished antibody reactivity while removal of only sialic acids or terminal fucoses improved the reactivity. The RBD produced in Lec3.2.8.1-cells, which generate carbohydrate structures devoid of sialic acids and with reduced fucose content, exhibited enhanced antibody reactivity verifying the importance of these specific monosaccharides. The results can be of importance for the design of future vaccine candidates, indicating that it might be possible to enhance the immunogenicity of recombinant viral proteins.

## Introduction

The adaptive immune response to SARS-COV-2 depends on T-cells that directs the immune responses and contributes to killing of infected cells, and on antibody producing B-cells (Rydyznski Moderbacher *et al*, 2020). Seroconversion has been detected in 93-99 % of patients with diagnosed SARS-CoV-2 infection, with disease severity correlating with antibody titres (Kellam & Barclay, 2020; Lou *et al*, 2020; Zhao *et al*, 2020). Neutralizing antibodies (NAb) are a key component in the response towards viruses, and an important aspect after immunization is whether the generated antibodies possess neutralizing capabilities (Plotkin & Plotkin, 2008). In the case of SARS-CoV-2 many neutralizing antibodies recognize the receptor binding motif (RBM) within the receptor binding domain (RBD) of the spike (S) protein (Tortorici *et al*, 2020), and execute their neutralizing capacity by sterically hindering viral binding to the angiotensin converting enzyme 2 (ACE2) receptor (Ju *et al*, 2020), or by keeping the RBD in its “down-conformation” (Tortorici *et al*., 2020). However, there are reports about neutralizing antibodies targeting epitopes also outside the RBM, such as 47D11 which binds to the conserved core of RBD (Wang *et al*, 2020), or S309, which recognizes a conserved epitope involving interactions with the fucose and other glycan moieties of the N343-glycan within the RBD (Pinto *et al*, 2020). Neutralizing antibodies have also been found to target the N-terminal domain (NTD) of the S protein. For example, the NAb 4A8 targets residues within the NTD, including the N147-glycosite, and neutralizes possibly by inhibiting conformational changes of the S protein (Chi *et al*, 2020). Thus, the glycosylation profile of the S protein appears to be important for antibody recognition and neutralization.

The S protein is inserted in the viral envelope as trimers, forming the characteristic “spikes” protruding from the viral surface. The S protein is cleaved by host proteases to form S1 and S2. The S1 domain contains the RBM and mediates binding to the ACE2 receptor, while fusion with the host cell membrane is mediated by S2 (Cavanagh, 1983; Delmas & Laude, 1990; Li *et al*, 2003). The S protein ectodomain contains 22 consensus sites for N-linked glycosylation (Asn-X-Ser/Thr where X is any amino acid except Pro). Most of the sites have been reported as glycosylated, carrying complex and high-mannose glycans for recombinant S proteins, expressed in cell culture (Allen *et al*, 2021; Sanda *et al*, 2021; Shajahan *et al*, 2020; Watanabe *et al*, 2020a). Although most consensus sites for N-linked glycosylation in the S protein appear to be occupied, the composition and structure of glycans at respective site appears highly variable (Allen *et al*., 2021; Watanabe *et al*., 2020a). The O-linked glycosylation pattern of the S protein is not entirely established, although the presence of several O-linked glycans have been identified within the RBD (Antonopoulos *et al*, 2021; Bagdonaite *et al*, 2021; Sanda *et al*., 2021; Shajahan *et al*., 2020).

The vaccines against SARS-CoV-2 induce antibodies that after immunization, target specific domains of the S protein. The vector based DNA vaccines and mRNA vaccines utilize the human glycosylation profile on the produced protein, while the glycosylation profile of protein-based sub-unit vaccines is dependent on the cell type used for production (Croset *et al*, 2012). As the glycosylation profile of the S protein may affect the antibody epitopes, leading to variability in the effectivity of the vaccine, the glycosylation of the target protein is an important issue to study.

In this work we have characterized the glycan content of a recombinant RBD protein expressed in three different mammalian cell lines and showed a diverse glycan composition at each site. The N- and O-linked glycans were stepwise modulated using enzymatic degradation. Serum samples from patients previously infected with SARS-CoV-2 were used to assess the impact of glycan composition on antibody reactivity. A glycan hot spot within the RBD was found to be essential for antibody reactivity. In addition, modulation of the glycan content revealed specific monosaccharides that were able to enhance the antibody reactivity.

## Results

### Glycosylation pattern of the recombinant RBD produced in CHO-S and HEK293F cells

Recombinant RBD, produced in HEK293F- and CHO-S-cells respectively, was subjected to nanoLC-MS/MS analysis. We defined the level of occupancy, composition, and structure of the N-linked and O-linked glycans present in the RBD. The HEK293F-produced RBD showed nearly complete occupancy for both N-linked sites (99.1 % and 100 %), while the CHO-S produced construct presented a partial occupancy of 93.3 % for site N331 and full occupancy for N343 (Fig. 1A and 1B). Complex type N-linked glycans were the most abundant structure in both cell lines, still a higher degree of glycans processed to complex type was associated with the HEK293F-cell line while oligomannose structures were relatively more abundant for the CHO-S-produced protein. The observed CHO-S-produced oligomannose glycans were different at the two sites with N331 displaying higher levels of oligomannose-6-phosphate glycans (Fig. 1A, 1B, and Table 1).

**Figure 1.**
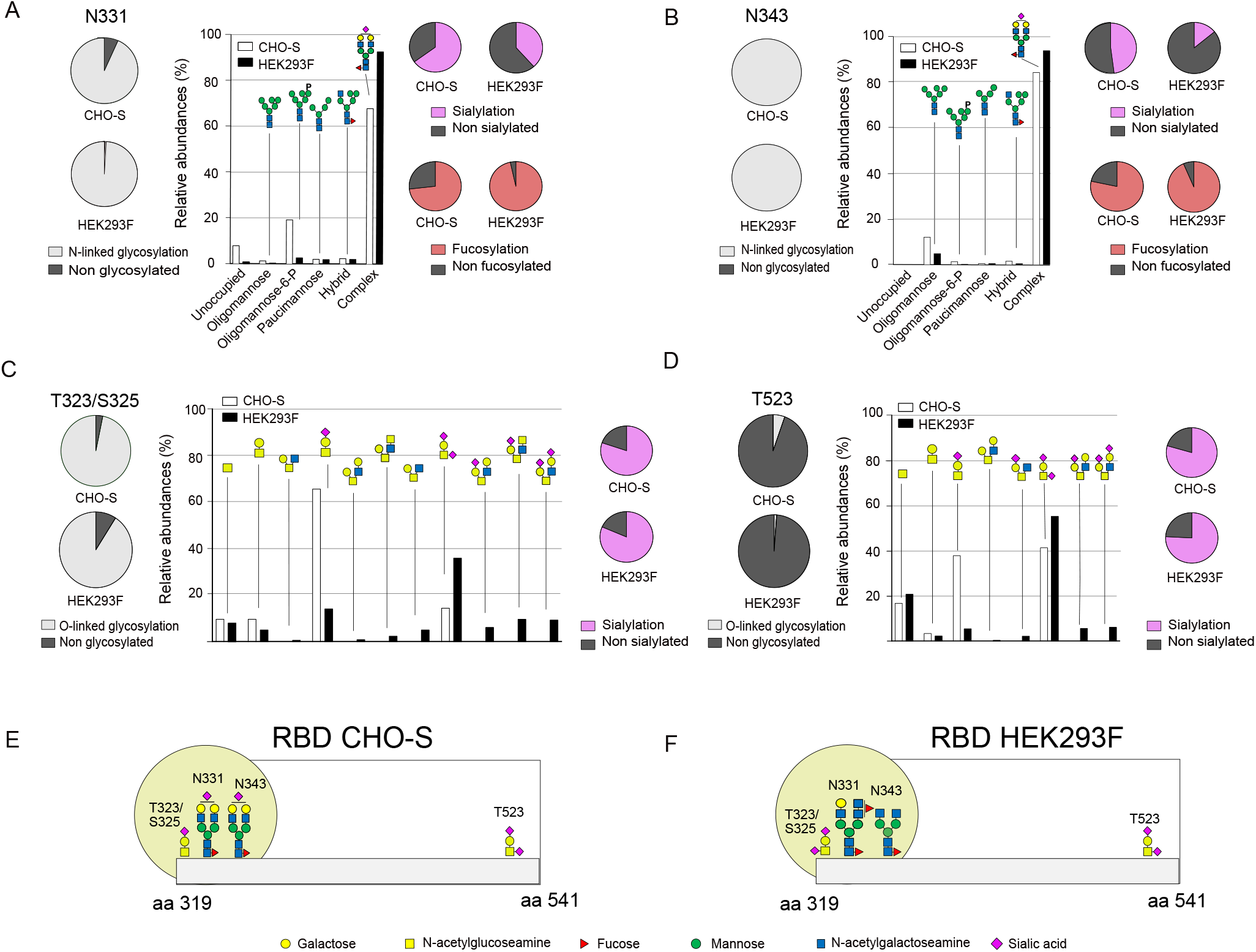
Schematic presentation of glycan distribution at the respective sites of the CHO-S- and HEK293F-produced RBD. A Degree of glycosylation, distribution between the glycan types, degree of sialylation and degree of fucosylation at site N331 as detected on CHO-S- and HEK293F produced RBD. B. Degree of glycosylation, distribution between the glycan types, degree of sialylation and degree of fucosylation at site N343 as detected on CHO-S- and HEK293F produced RBD. C. Degree of glycosylation, distribution between glycan composition and degree of sialylation at site T323/S325 as detected on CHO-S and HEK293F-produced RBD. D. Degree of glycosylation, distribution between glycan composition and degree of sialylation at site T523 as detected on CHO-S and HEK293F-produced RBD. E. Glycosylation of recombinant RBD produced in CHO-S cells with the most prevalent glycans drawn at the respective site. Yellow circle highlights the glycan hotspot. F. Glycosylation of recombinant RBD produced in HEK293F cells with the most prevalent glycans drawn at the respective site. Yellow circle highlights the glycan hotspot.

**Table 1.** Percentage distribution of glycan types at site N331 and N343 when produced in CHO-S, HEK293F and Lec3.2.8.1-cells.

|  | N331 |  |  | N343 |  |  |
| --- | --- | --- | --- | --- | --- | --- |
|  | CHO-S | HEK293F | Lec3.2.8.1 | CHO-S | HEK293F | Lec3.2.8.1 |
| Unoccupied | 7.8 | 0.9 | 2.5 | 0.0 | 0.0 | 0.0 |
| Oligomannose | 1.3 | 0.4 | 76.1 | 12.1 | 4.9 | 79.8 |
| Oligomannose-6-P | 19.1 | 2.7 | 10.5 | 1.4 | 0.2 | 2.3 |
| Paucimannose | 2.1 | 2.0 | 8.1 | 0.6 | 0.6 | 12.4 |
| Hybrid | 2.5 | 2.1 | 2.2 | 1.6 | 0.5 | 5.5 |
| Complex | 67.3 | 91.8 | 0.6 | 84.3 | 93.8 | 0.0 |

Among the complex type N-linked glycans, biantennary structures were most frequently found at both positions (Table EV2). Despite similar glycan compositions in both cell lines, the fragment spectra evaluation identified prominent differences for glycans produced in CHO-S and HEK293F-cells. The major difference was the prominent LacDiNAc-containing structures in HEK293F-produced RBD, while those were absent in the CHO-S-produced protein (Table EV2).

Within a given cell line, the frequency of fucose residues was similar for both sites, while the overall fucosylation was higher for HEK293F compared to CHO-S (Fig. 1A, 1B, and Table 2). Not only the total fucosylation level, but also the degree of fucosylation (number of fucose residues per glycan) differ between the cell lines (Table EV3). CHO-S cells predominantly produced monofucosylated structures with the fucose placed at the core, as based on the fragment ion analysis. Multiple fucosylation, with up to four fucose residues per glycan, was observed for RBD produced in the HEK293F cell line. The attachment of fucose to LacNAc and LacDiNAc was observed based on the fragment spectra evaluation.

**Table 2.** Percentage of detected glycans carrying at least one fucose or sialic acid at site N331 and N343 when produced in CHO-S, HEK293F and Lec3.2.8.1-cells.

|  | N331 |  |  | N343 |  |  |
| --- | --- | --- | --- | --- | --- | --- |
|  | CHO-S | HEK293F | Lec3.2.8.1 | CHO-S | HEK293F | Lec3.2.8.1 |
| Fucose* | 73.2 | 96.6 | 31.4 | 78.4 | 93.5 | 6.3 |
| Sialic acid** | 65.0 | 38.4 | 0.0 | 47.9 | 14.1 | 0.0 |

\* The percentage is calculated in relation of total amount of all observed glycosylated forms, unoccupied peptides were excluded when performing the calculations.
\*\* The percentage is calculated in relation of total amount of all observed hybrid- and complex-type glycoforms, the unoccupied peptides and all oligomannose glycoforms were excluded from the calculations.

In contrast to fucosylation, the sialylation level was lower for HEK293F compared to CHO-S (Fig. 1A, 1B, and Table 2). Also, the degree of sialylation (number of sialic acid residues per glycan) differed between the cell lines (Table EV4). CHO-S-cells produced multiple sialylated forms in contrast to HEK293F, where mainly mono-sialylated structures were observed. The lower sialylation level in glycans produced by HEK293F is likely a result of the extensive fucosylation in this cell type. In both cell lines the major N-glycan type carrying sialic acid was complex type glycans and a difference between the sites were noted, with N-linked glycans at position N331 displaying higher sialylation levels.

The O-linked glycans were similar for the two cell types (Fig. 1C, 1D, and Table EV5). The O-linked glycan close to the N-terminal domain of the RBD could not be defined to a single amino acid, due to the absence of fragment ions between the two adjacent potential sites, and thus could be placed either at amino acid position T323 or S325. The site T323/S325 was glycosylated to a high degree (97 % and 91 %, for CHO-S and HEK293F respectively), while T523 was scarcely decorated and mainly remained non-glycosylated in both CHO-S- and HEK293F-produced RBD (5 % and 1 %, respectively). Comparison of O-linked glycans at the individual sites revealed more extensive processing and branching in the HEK293F produced protein while the CHO-S produced O-linked glycans almost exclusively consisted of core 1 structures (Fig. 1C and 1D). The degree of sialylated structures at site T323/S325 were similar between the cell lines (80 % and 82 % for CHO-S and HEK293F, respectively), while the degree of monosialylated structures (66 % and 35 %, respectively) and disialylated structures (15 % and 46 %, respectively) differed. Similarly, the frequency of sialylated structures at site T523 was similar between CHO-S (80 %) and HEK293F (79 %) cells. The degree of monosialylation was higher in the CHO-S produced RBD, as compared to the HEK293F produced protein (38 % and 14 %, respectively), while a higher degree of disialylation was seen on the HEK293F produced RBD (42 % and 62 %, respectively) (Fig. 1C and 1D).

In summary, RBD produced in CHO-S cells carried O-linked glycans at two positions although only position T323/S325 appeared to be glycosylated with high frequency. The main type of O-linked glycan found at this position was a core 1 structure with a single sialic acid at the distal galactose (Fig. 1E). This RBD protein also carried two N-linked glycans at position N331 and N343. The predominant type of N-linked glycan was the biantennary complex type, although many variants of complex type glycans were found. RBD produced in HEK293F cells predominantly carried core 1 O-linked glycans with two sialic acids, one attached to the distal galactose and one to the innermost GalNAc residue. The N-linked glycans on the HEK293F RBD were almost exclusively of complex type with a high degree of fucosylation (Fig. 1F).

### Evaluation of convalescent sera from Covid-19 patients

Serum samples were collected from 24 individuals previously infected with SARS-CoV-2 as determined by a PCR-positive nasopharyngeal sample. Blood samples were collected 25 – 100 days following positive diagnosis. All sera were characterized with respect to anti-RBD IgG levels and the capability to neutralize a DE-Gbg20 strain of SARS-CoV-2 grown in VERO-cells. Based on the neutralization capability the sera were divided to three groups: non-neutralizing (NT negative, n=7), weakly neutralizing (NT titre 3-6, n=7) and highly neutralizing (NT titre 48-96, n=10) (Fig. 2 and Table EV6). High neutralization capability correlated well with high levels of IgG targeting the RBD, as all highly neutralizing sera also were anti-RBD IgG positive while six of the seven serum samples in the weakly neutralizing group were anti-RBD IgG negative. Interestingly, four out of seven serum samples in the NT negative group were anti-RBD IgG positive.

**Figure 2.**
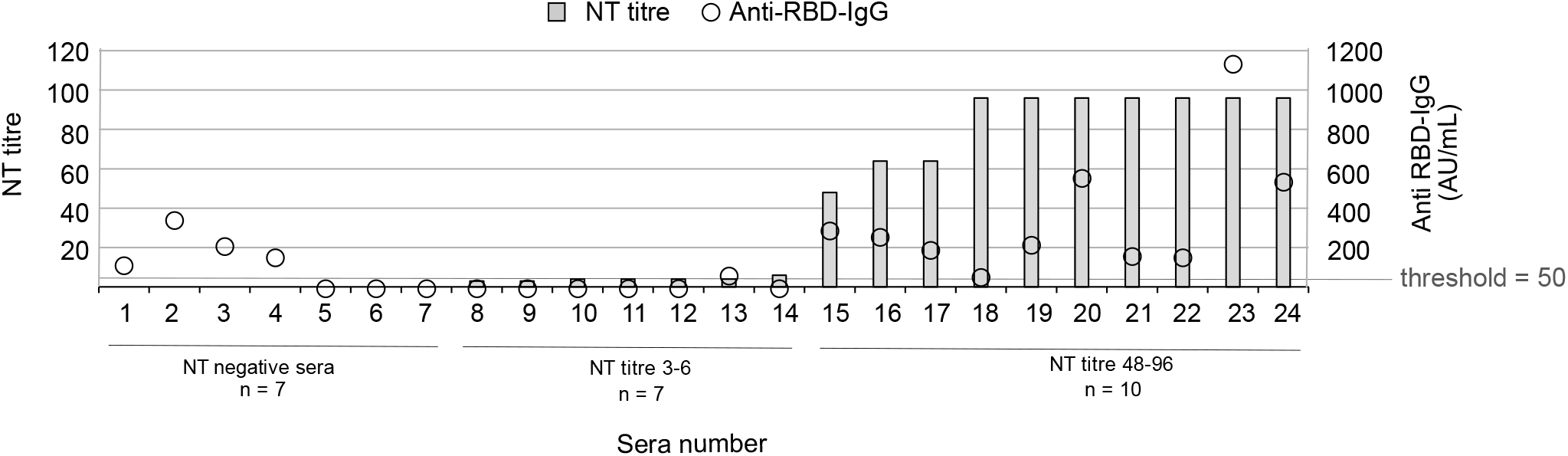
NT-titre (left y-axis, grey bars) and anti-RBD IgG-levels (right y-axis, transparent circles) for the 24 characterized serum samples. The sera were divided into three groups based on the neutralizing capability: non-neutralizing (NT-negative, n=7), weakly neutralizing (NT titre 3-6, n=7) and highly neutralizing (NT titre 48-96, n=10). Anti RBD-IgG value ≥ 50 AU/mL is considered positive.

### Impact of glycan structures on antibody reactivity against RBD

To assess the impact of the different types of glycan structures found within the RBD we removed the N-linked, the O-linked or a combination of both glycans using enzymatic treatment. Removal of glycans was verified by a size shift on an SDS-page gel, visualized by silver staining (Fig. EV1A). The effect of glycan removal on the antibody reactivity against the recombinant RBD was tested using the defined serum samples described above. RBD produced in CHO-S-cells elicited a strong reactivity to the highly neutralizing sera, with reduced reactivity following removal of N-linked glycans, O-linked glycans or a combination of both (Fig. 3A). The highly neutralizing sera showed reduced reactivity against the RBD with removed N-linked glycans and the RBD lacking both N- and O-linked glycans produced in HEK293F-cells, while no effect on reactivity against the RBD lacking only O-linked glycans was observed (Fig. 3B). Weakly neutralizing sera did not show any reactivity to the CHO-S- or the HEK293F-produced RBD regardless of the glycosylation profile (Fig. EV2A and EV2C). The non-neutralizing serum samples displayed low reactivity against all recombinant RBD with only a minor difference depending on glycosylation status (Fig. EV2B and EV2D). The intensity of the reactivity of individual serum samples against the recombinant RBD correlated well with the anti-RBD IgG levels detected in each serum (Fig. EV3).

**Figure 3.**
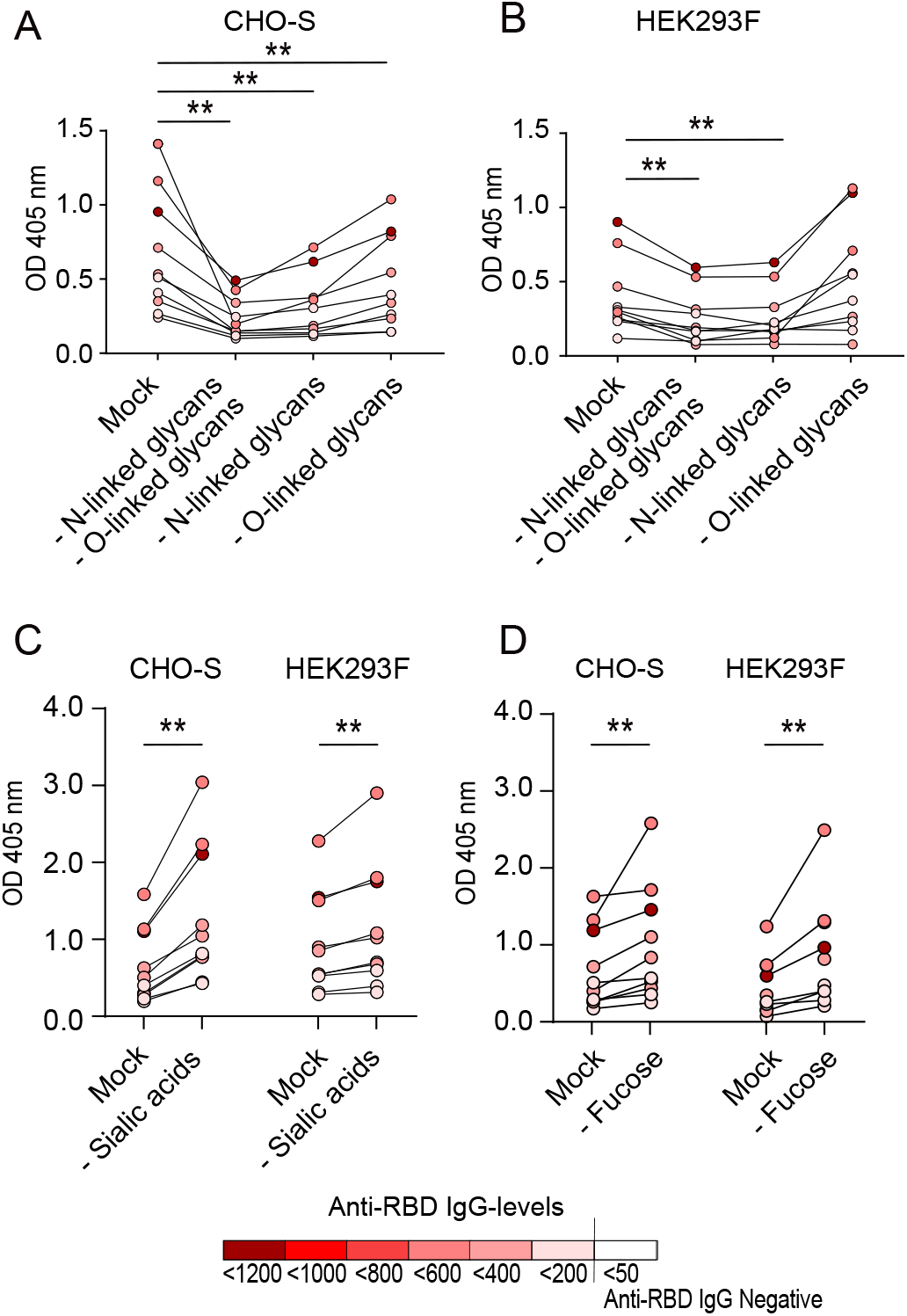
Reactivity of highly neutralizing sera (NT titre 48-96, n=10) against the fully glycosylated RBD (mock treated) and against deglycosylated RBD produced in CHO-S and HEK293F-cells. A. Removal of both N-linked and O-linked glycans, removal of N-linked glycans alone, or removal of O-linked glycans alone from RBD produced in CHO-S-cells. B. Removal of both N-linked and O-linked glycans, removal of N-linked glycans alone, or removal of O-linked glycans alone from RBD produced in HEK293F-cells. C. Removal of sialic acids alone from RBD produced in CHO-S and HEK293F-cells. D. Removal of fucose alone from RBD produced in CHO-S and HEK293F-cells. Data information: Dark red colour symbolizes a serum with high levels of anti-RBD IgG, white colour indicates anti-RBD IgG-negative serum (<50 AU/mL). Statistical analysis was performed with the Wilcoxon matched-pair signed rank test, ** = p<0.001.

To further assess the impact of specific glycan residues on antibody reactivity, sialic acids and fucose groups were enzymatically removed from the CHO-S and HEK293F-produced RBD. SDS-page gel electrophoresis and subsequent silver stain was used to confirm the removal of sialic acids or fucose groups. Small, but distinct, size shifts were evident after the enzymatic treatments, indicating successful removal of each glycan species (Fig. EV1B and EV1C). Removal of sialic acids from the CHO-S-produced RBD resulted in a significant increase in the serum reactivity, as compared to fully glycosylated RBD. A similar effect was seen following removal of sialic acids from the HEK293F-produced RBD, however this difference was less prominent (Fig. 3C). Removal of fucose groups also resulted in a significant increase in serum reactivity for both the CHO-S- and HEK293F-produced RBD, with the HEK293F-produced construct showing a more prominent increase (Fig. 3D).

To confirm the impact of sialic acids and fucose groups on the antibody reactivity, the RBD was produced in Lec3.2.8.1-cells deficient in synthesis of complex type glycans. Highly neutralizing serum samples showed a significantly higher reactivity against RBD produced in Lec3.2.8.1-cells, as compared to the CHO-S- or HEK293F-produced RBD constructs (Fig. 4A). As expected, enzymatic removal of sialic acids and fucose from the Lec3.2.8.1 produced RBD, did not confer any detectable size shift on an SDS-page gel (Fig. EV1D) or a change in antibody reactivity by highly neutralizing sera (Fig. 4B).

**Figure 4.**
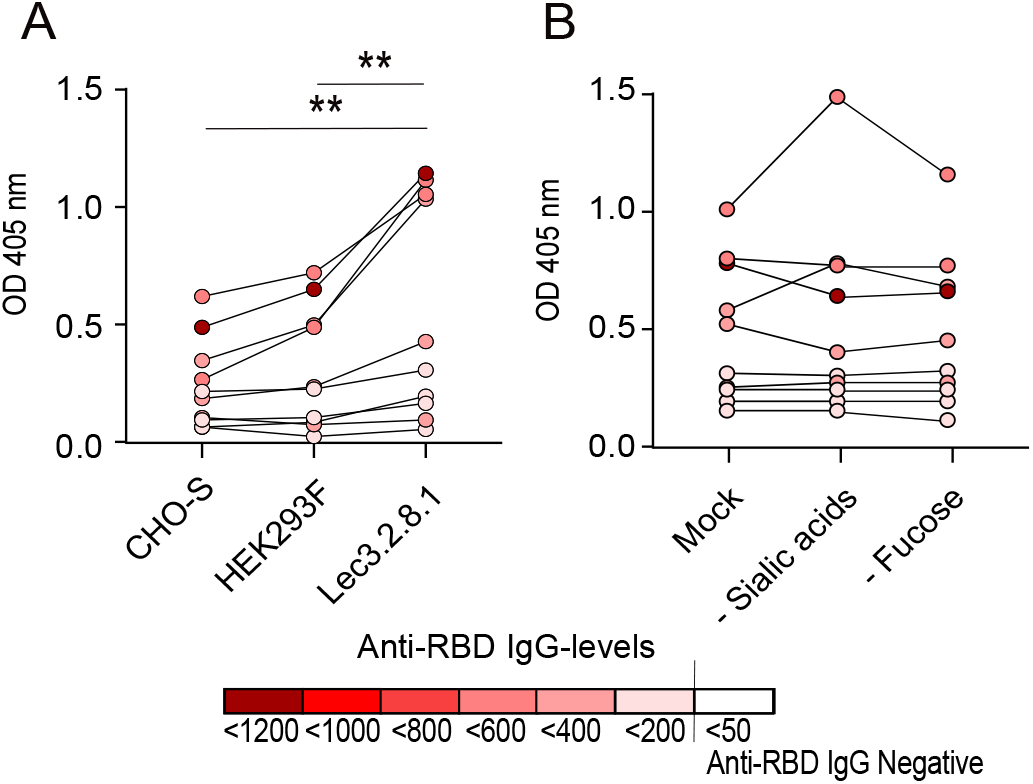
Antibody reactivity of highly neutralizing sera (NT titre 48-96, n=10). A. Reactivity against the fully glycosylated RBD (untreated) expressed in CHO-S, HEK293F and Lec3.2.8.1-cells. B. Reactivity against the fully glycosylated RBD (mock treated) produced in Lec-3.2.8.1-cells, or against the Lec3.2.8.1 produced RBD following enzymatic removal of sialic acids and fucose. Data information: Dark red colour symbolizes a serum with high levels of anti-RBD IgG, white colour indicates anti-RBD IgG-negative serum (<50 AU/mL). Statisitical analysis was performed with Wilcoxon matched-pair signed rank test, ** = p<0.001.

### Glycosylation of recombinant RBD produced in Lec3.2.8.1-cells

In order to verify that the recombinant RBD produced in Lec3.2.8.1-cells lacked complex type glycans, it was subjected to nanoLC-MS/MS analysis. Both N-linked sites were found to be glycosylated to a high degree (98 % and 88 %, respectively). The structural distribution was similar between the sites, with high-mannose as the dominating glycan type (Table 1). No sialic acid or end-fucose were found, however 6 % of the structures at site N331 and 31 % of the structures at site N343 carried core fucose (Table 2). Position T323/S325 was frequently (98 %) decorated with an O-linked glycan, while site T523 were more sparsely decorated (15 %). A single HexNAc was the most frequent structure at both O-linked sites (Table EV5).

## Discussion

Virus that infects humans do not carry their own glycosyltransferases, but instead relies on the enzymes of the host cell in the processing of their glycans. This often results in a viral glycosylation profile with traits similar to the host cell, that does not stimulate a strong immune response. Thus, in order to circumvent the antibody responses, many viruses utilizes the host cell glycosylation machinery to cover B-cell epitopes with a dense network of N-linked glycans, which due to physical hindrance shield the epitopes and prevent binding by neutralizing antibodies (Grant *et al*, 2020). With a few exceptions, N-linked glycans alone rarely acts as antibody epitopes (Raska *et al*, 2010), but the opposite effect has been observed with small O-linked glycans. Olofsson et al. showed that 70 % of tested sera against herpes simplex virus (HSV) type 2 contains antibodies targeting a peptide decorated with a single O-linked GalNAc residue. Removal of the glycan moiety diminished this response (Olofsson *et al*, 2016). Similarly, our group has previously shown that a single GalNAc-residue added to a naked peptide can alter the antibody binding towards specific domains of the glycoprotein E from the varicella zoster virus (VZV) (Nordén *et al*, 2019).

It is established that viral glycoproteins are heterogeneously glycosylated when they are expressed either as recombinant proteins in cell culture or after natural infection in cells (Brun *et al*, 2021; Nordén *et al*, 2015; Nordén *et al*., 2019). This could imply that the antibody response to a viral glycoprotein is more diverse than previously thought. Hence, serum from infected individuals contain a polyclonal antibody pool which could recognize multiple epitopes and various glycoforms can constitute parts of these epitopes. The S protein of SARS-CoV-2 is highly glycosylated, with 17 to 22 previously identified sites carrying N-linked glycosylation that can shield B cell epitopes (Allen *et al*., 2021; Antonopoulos *et al*.,2021; Sanda *et al*., 2021). Of these, 2 N-linked glycans are present in the RBD, and Yang et al. identified as many as 10 O-linked glycans in this region, although most of them appeared to be of low abundance and their biological significance is therefore uncertain (Yang *et al*, 2020). We demonstrate that enzymatic removal of all N-linked and/or O-linked glycans resulted in decreased antibody reactivity of the recombinant SARS-CoV-2 RBD produced in both CHO-S and HEK293F cells. This could indicate (i) that the glycans of the RBD constitute parts of antibody epitopes, (ii) that glycosylation is required for proper protein folding and in maintaining the protein conformation (Shental-Bechor & Levy, 2008) or (iii) a combination of both.

The strong immunoreactivity of RBD, as verified by the identification of a multitude of Nabs targeting this domain (Pinto *et al*., 2020; Tortorici *et al*., 2020; Wang *et al*., 2020), could be explained by the presence of multiple structural epitopes. If large glycan moieties are lacking, as after enzymatic removal, the folding of the protein could be compromised and thereby also transform structural epitopes which would render them inaccessible to antibodies present in the serum samples. No linear B-cell epitopes have been identified within the RBD (Li *et al*, 2021). However, screening for B cell epitopes is performed with synthetic peptides lacking glycans and would then not be able to identify epitopes that are dependent on the presence of N-or O-linked glycans.

We performed a structural screening of the glycan profile of recombinant RBD. Overall, our screening is in line with previous studies (Shajahan *et al*., 2020; Watanabe *et al*, 2020b), but we found significant differences in the amount of sialic acid and fucose content when comparing the RBD produced in CHO-S cells and HEK293F cells respectively. Interestingly, the CHO-S-produced RBD also presented with a high degree of mannose-6-phospate (M-6-P). This structure has previously been observed in the SARS-CoV-2 spike protein when expressed in cell lines but also when isolated from intact viral particles (Brun *et al*., 2021; Gstöttner *et al*, 2021). Mannose-6-phosphate is recognized by the M-6-P receptor present in the trans-Golgi compartment and it directs tagged proteins to late endosomes/lysosomes. Lysosomal egress dependent on M-6-P has been described for both HSV (26) and VZV (27). Also, SARS-CoV-2 egress mediated by lysosomes has been proposed (Ghosh *et al*, 2020). However, we observed only a minor fraction of the peptides carrying M-6-P and to what extent they potentially could contribute to viral particle egress remains to be clarified.

While each glycosite on the recombinant RBD was glycosylated at a similar frequency independent of production cell line, RBD produced in HEK293F had a higher degree of fucosylation compared to CHO-S. Selective removal of the fucose groups resulted in a significantly increased antibody reactivity. While abundant fucosylation was a trait of the HEK293F produced construct, RBD produced in CHO-S cells had a larger content of sialic acid moieties. Selective removal of sialic acids enhanced antibody reactivity for both constructs, but the effect was more prominent for the CHO-S construct. The use of the Lec3.2.8.1-cell line resulted in RBD with a glycosylation profile completely deficient in sialic acids and end-fucose. Consequently, the antibody reactivity towards the Lec3.2.8.1-produced RBD was enhanced and additional removal of the core-fucose did not result in any change in the antibody reactivity. Altogether, these results points to an important function of specific terminal-sugar residues in the antibody reactivity against glycosylated viral antigens and suggests that that core-fucosylation is of minor importance, despite the report of an NAb that specifically interacts with the core fucose of the N-linked glycan situated on position N343 (Pinto *et al*., 2020).

In line with our findings that removal of sialic acids leads to increased antibody reactivity, the non-sialylated glycan structures of yeast-cell produced proteins could possibly be part of the explanation of the highly efficient yeast-produced vaccines against HBV (Doering *et al*, 2015; Ho *et al*, 2020). This suggests that it is possible to optimize recombinantly expressed RBD or S proteins in order to generate effective vaccine candidates. However, important to note is the possibility that immunization with a recombinant expressed subunit vaccine directs the humoral immune response towards B-cell epitopes with species-specific glycosylation profiles. This can possibly result in skewed immunodominance, directing the antibody response towards epitopes that are not exposed after a natural infection with the virus, resulting in disturbed efficiency of the vaccine (Abbott & Crotty, 2020). The data presented in this work confirms the necessity of correct glycosylation, and shows that also small differences in the glycosylation profile of a viral antigen can have a large impact on the reactivity by antibodies generated after a natural infection with SARS-CoV-2. A conscious decision regarding the glycosylation traits of the production cell line could hence affect the antibody response triggered by a recombinant protein. We suggest the glycosylation characteristics should be considered during the production of recombinant vaccines towards SARS-CoV-2 but also other enveloped viruses which carry glycoproteins.

## Material and methods

### Expression of recombinant S protein constructs

The receptor-binding domain of the SARS-CoV-2 spike protein (amino acids 319-541) was produced in three cell lines using an expression vector obtained through BEI Resources, NIAID, NIH, which is vector pCAGGS containing the SARS-CoV-2, Wuhan-Hu-1 spike glycoprotein gene RBD with C-terminal Hexa-Histidine tag (NR-52309) (Table EV1). CHO-S cells (Cat nr R80007, Thermo Fisher Scientific, Waltham, MA) were adapted to grow in suspension in FectoCHO medium (Polyplus transfection, Illkirch-Graffenstaden, France) at 37 °C in 5 % CO_2_ in Optimum Growth™ flasks (Thomson instrument company, Oceanside, CA) at 130 rpm in a Multitron 4 incubator (Infors, Bottmingen, Schweiz). Lec3.2.8.1 cells (a mutated CHO cell line kindly received from Prof. P Stanley (Chen & Stanley, 2003)) were cultured under the same conditions. The HEK293 derivate HEK293F cell line (Cat nr R79007, Thermo Fisher Scientific) were cultured in Freestyle 293 medium. Cells were transfected at 2 × 10^6^ cells/mL using FectoPro transfection reagent (Polyplus transfection). The temperature was reduced to 32 °C (Lec3.2.8.1) or 31 °C (CHO-S) 4 hours post transfection, while transfected HEK293F cells were kept at 37 °C. Protein-containing culture supernatants (800 mL – 1 L) were harvested when cell viability was below 80 %, which was after 168 hours (CHO-S), 74 hours (Lec3.2.8.1) or 90 hours (HEK293F), filtered using Polydisc AS 0.45 μm (Whatman, Maidstone, UK) and loaded onto a 5 mL HisExcel column (Cytiva, Marlborough, MA). After sample loading, the column was washed with 20 mM sodium phosphate, 0.5 M NaCl and 30 mM imidazole before elution of the protein using the same buffer with 500 mM imidazole (Lec3.2.8.1-produced RBD) or 300 mM imidazole (CHO-S and HEK293F-produced RBD). Pooled fractions were concentrated using 10 kDa Vivaspin concentrators (MWCO 10 kDa, Sartorius, Göttingen, Germany), passed over a HiPrep 26/10 desalting column (Cytiva) in phosphate-buffered saline and finally concentrated again. The Lec3.2.8.1-produced RBD was further purified by gel filtration using a Superose 200 Increase 16/300 GL column (Cytiva) in phosphate-buffered saline. Integrity and purity of the different RBD preparations were checked by SDS-PAGE and Western blot.

### Sample preparation prior to assessment of position and structure of glycans

The purified RBD preparations from CHO-S, HEK293F and Lec3.2.8.1 (20 μg each) were diluted with digestion buffer (DB), 1 % sodium deoxycholate (SDC) in 50 mM triethylammonium bicarbonate (TEAB) pH 8.0 (Sigma Aldrich, St. Louis, MO), to give protein concentrations of 0.5 μg/μL. The RBD preparations were reduced with 4.5 mM dithiothreitol (DTT) at 56 °C for 30 min and alkylated with 9 mM 2-iodoacetamide (IAM) in the dark for 30 min at room temperature (RT). The alkylation reactions were then quenched by incubation with DTT (9 mM final concentration) for 15 minutes at RT. Additional 20 μL of DB was added prior to the proteolytic digest with Pierce™ MS grade trypsin and Glu-C (overnight at 37 °C, 0.2 μg and 0.3 μg, respectively). The digested samples were purified using High Protein and Peptide Recovery Detergent Removal Spin Column (Thermo Fisher Scientific) according to the manufacturer instructions. SDC was removed by acidification with 10% trifluoroacetic acid (TFA) and subsequent centrifugation.

The supernatants were further purified using Pierce peptide desalting spin columns (Thermo Fisher Scientific) according to the manufacturer’s instructions. Each of the purified RBD preparations was divided into 3 parts: (1) 7.5 μg for nanoLC-MS/MS analysis, (2) 7.5 μg for neuraminidase treatment, (3) 5 μg for PNGaseF treatment.

For sialic acid removal, RBD preparations were incubated with 1 μl Sialidase A (GK80040, Agilent, Santa Clara, CA) in 50 μL of provided buffer, overnight at 37 °C. For N-glycans removal, samples were dissolved in 50 μL of 50mM TEAB and treated with 1 μL of recombinant PNGaseF (Promega, Madison, WI) overnight at 37 °C. All preparations were desalted using Pierce Peptide Desalting Spin Columns (Thermo Fisher Scientific) prior to NanoLC-MS/MS analysis.

### NanoLC-MS/MS analysis of recombinant RBD

The RBD proteolytic preparations were analyzed on a QExactive HF mass spectrometer interfaced with Easy-nLC1200 liquid chromatography system (Thermo Fisher Scientific). Peptides were trapped on an Acclaim Pepmap 100 C18 trap column (100 μm × 2 cm, particle size 5 μm, Thermo Fischer Scientific), and separated on an in-house packed analytical column (75 μm × 300 mm, particle size 3 μm, Reprosil-Pur C18, Dr. Maisch) using a gradient from 7 % to 50 % B over 75 min, followed by an increase to 100 % B for 5 min at a flow of 300 nL/min, where Solvent A was 0.2 % formic acid (FA) and solvent B was 80 % acetonitrile (ACN) in 0.2 % FA. The precursor ion mass spectra were acquired in either 600-2000 m/z or 375-1500 m/z ranges at a resolution of 120 000. For NanoLC-MS/MS analysis, the instrument operated in data-dependent mode with the 10 most intense ions with charge states 2 to 5 being selected for fragmentation using higher-energy collision dissociation (HCD). The isolation window was set to 3 m/z and dynamic exclusion to 20 s. MS/MS spectra were recorded at a resolution of 30 000 with maximum injection time set to 110 ms. To facilitate glycosylated peptide characterization, multiple injections were acquired with precursors detection in 600-2000 m/z range and different settings for the normalized HCD energies of 22, 28 and 34.

### Glycan database search and data processing

The acquired data were analyzed using Proteome Discoverer version 2.4 (Thermo Fisher Scientific). Database searches were performed with either Byonic (Protein Metrics, Cupertino, CA) or Sequest as search engines. To evaluate the protein preparations purity, the data were initially searched against custom database consisting of Uniprot_Chinese hamster_CHO-K1 cell line database (24147 proteins), SwissProt_human database (20342 proteins) and the sequence of expressed RBD protein. For later searches aimed to identify the available glycoforms, the raw data acquired with different HCD energies were searched with Proteome Discoverer/Byonic, with Minora Feature Detector node, against the single RBD protein sequence. Precursor mass tolerance was set to 10 ppm and fragment mass tolerance to 30 ppm. Proteolytic peptides with up to 2 missed cleavages (combined Trypsin Glu-C cleavage sites) were accepted together with variable modification of methionine oxidation and fixed cysteine alkylation. Several different N-glycan databases were used during the data processing.

The initial N-glycan database contained 227 glycan compositions, where 224 were COVID-19 associated N-glycan compositions reported in GlyConnect Compozitor version: 1.0.0 at SIB Swiss Institute of Bioinformatics | Expasy site plus 3 additional mannose-phosphate containing compositions. This database was used for the analysis of neuraminidase treated samples. The 45 curated non-sialylated compositions, retrieved from the analysis of neuraminidase treated preparation, were used to create a new glycan database consisting of 153 glycan compositions for the follow-up analysis of native preparations. An O-glycan database consisted of 6 reported in GlyConnect Compozitor COVID-19 O-glycan compositions and was used for the analysis of PNGaseF treated samples.

All glycopeptide identifications were manually evaluated prior to the final assignment of the observed glycosylation forms. The data, acquired with the normalized HCD energy of 22, were used for oxonium ion evaluation to suggest glycan structures for the observed compositions. The extracted ion chromatogram (EIC) peak intensities of the observed glycoforms were used to calculate their relative abundances. The relative abundances were calculated using average EIC values from the multiple injections and are expressed as percent of the total signal for all modified and non-modified forms.

### Deglycosylation of recombinant RBD using glycosidase treatment

Removal of N-linked glycosylation was performed using PNGaseF (New England Biolabs, Ipswich, USA) at a concentration of 125 U/μg protein. Removal of both N-linked and O-linked glycans was performed by incubation with five different glycosidases: PNGaseF (New England Biolabs, 125 U/μg protein), O-glycosidase (New England Biolabs, 20 000 U/μg protein), α2-3,6,8 neuraminidase (New England Biolabs, 25 U/μg protein) and α-N-acetyl-galactosaminidase (New England Biolabs, 10 U/μg protein). Removal of sialic acids was performed using the α2-3,6,8 neuraminidase (New England Biolabs, 50 U/μg protein). Removal of fucose was performed using α1-2,4,5,6 fucosidase O (New England Biolabs, 2 U/μg protein), and α1-3,4 fucosidase (New England Biolabs, 4 U/μg protein). All enzymatic reactions were performed as a 1-step reaction with 1x Glycobuffer 2 (New England Biolabs), 10 μg RBD produced in CHO-S-, HEK293F-, or Lec3.2.8.1-cells and incubation at 37 °C for 24 hours. As heat-treated controls, peptides were incubated at 37 °C for 24 hours, but without additional enzymes.

### Gel electrophoresis and in-gel staining of glycosidase treated recombinant RBD

To control efficiency of the enzymatic treatment 5 μg of the enzyme-treated products or controls were run on a NuPage™ 4-12 % Bis-Tris gel (Invitrogen, Carlsbad, USA) at 100 V for 60 minutes using the EI9001-XCELL II Mini Cell (Novex, San Diego, CA) together with the Powerease 500 (Novex) and subsequently stained with the SilverQuest™ Stain kit (Invitrogen, Carlsbad, USA) according to the instructions from the manufacturer.

### Levels of human anti-SARS-CoV-2 IgG antibodies in convalescent serum samples

Serum samples from SARS-CoV-2 convalescent individuals (n=24) were obtained from the department of Clinical Microbiology, Sahlgrenska University Hospital, Gothenburg, Sweden. Samples were collected between 06-03-2020 and 08-27-2020, 25 – 100 days following a positive PCR-test. Serum was stored at −80 °C until use. The SARS-CoV-2 IgG II Quant assay is a chemiluminescent microparticle immunoassay (CMIA) used for quantitative determination of IgG antibodies to SARS-CoV-2 in human serum and plasma on the ARCHITECT System (Abbott Laboratories, Chicago, IL). The assay measures IgG binding to the RBD of the S-protein. IgG concentrations ≥ 50 antibody units (AU)/mL were defined as positive.

### Viral CPE neutralization assay

The titre of neutralizing antibodies against SARS-CoV-2 in the patient sera was determined against the DE-Gbg20 viral strain (NCBI GenBank ID: MW092768) at a titre of 10^−6^. 50 % tissue culture infectious dose (TCID50) assay was performed as defined by Reed and Muench (Reed & Muench, 1938). All sera were heat inactivated at 56 °C for 30 minutes before 2-fold serial dilution in serum-free DMEM with 100TCID50 DEGbg20 followed by incubation for 2 hours at 37 °C. The virus-antibody mixture was added to a monolayer of VERO CCL-81 cells grown in 96 well-plates in DMEM supplemented with 2 % penicillin-streptomycin and 2 % foetal calf serum. The plates were incubated for 72 hours at 37 °C with 5 % CO_2_. The neutralizing titre for each serum was defined at the highest serum dilution at which 50 % of the added virus was neutralized.

### Anti-SARS-CoV-2 antibody reactivity assay

The antibody reactivity towards glycosidase treated proteins were assessed using an enzyme-linked immunosorbent assay (ELISA). Briefly, Nunc Maxisorp™ 96-well plates (Thermo Fischer Scientific) were coated with 0.1 μg glycosidase treated peptides or heat-treated controls diluted in carbonate buffer (pH 9.6). Coating was performed over night at 4 °C followed by washing three times with 0.05 % tween20 in phosphate buffered saline (PBS).

The plates were blocked in 2 % milk for 30 minutes at room temperature prior to addition of sera (diluted 1:100 in 1 % milk in PBS with 0.05 % tween20) and 1.5 hours incubation at 37 °C. The plates were washed three times before addition of alkaline phosphatase-conjugated Goat anti-human IgG (Jackson ImmunoResearch, Cambridgeshire, UK) diluted 1:1000 in 1 % milk in PBS with 0.05 % tween. After 1.5 h incubation at 37 °C the plate was washed six times and 1 mg/mL p-nitrophenylphosphate (Medicago, Danmarks-Berga, Sweden) in Diethanolamine Substrate buffer was added. The plates were incubated in the dark for 30 minutes before spectrophotometric measurement at 405 nm.

### Statistics

For the comparison of antibody reactivity as determined by ELISA the Wilcoxon matched-pair signed rank test was used. The comparison between anti-RBD IgG levels and antibody reactivity towards recombinant RBD was done with Pearson correlation coefficient, assuming normal distribution. All statistical analyses were performed using the Graphpad Prism software version 9.3.1 (GraphPad Software Inc, San Diego, CA, USA).

### Ethical statement

The study was approved by the ethical review board in Gothenburg (Dnr: 2021-02252).

## Acknowledgements

We thank Rickard Lymer, Mikael Andersson, and Vijay Kumar Nallani at the Mammalian Protein Expression Core Facilities at Gothenburg University for skilful production of recombinant constructs and Sigvard Olofsson for reviewing of the manuscript. We also thank The Swedish National Infrastructure for Biological Mass Spectrometry (BioMS) for financial support of glycoproteomic studies at the Proteomics Core Facility, University of Gothenburg. The study was funded by Sweden’s innovation agency Vinnova, (2020-03108).

## Author contributions

Conceptualization: RN; Formal analysis: ES, EM, KN; Funding acquisition: RN; Investigation; ES, EM, KN; Methodology: ES, EM, MB; Project administration: ES; Resources: KN, MB, JL; Supervision: EM, RN; Validation: ES; Visualization: ES; Writing – original draft: ES, RN; Writing – review and editing: ES, EM, KN, MB, JL, RN.

## The paper explained

### Problem

The SARS-CoV-2 spike protein contains carbohydrate structures and several studies have showed a large diversity in their composition. This is especially true for recombinant spike proteins, where the production cell line impacts the resulting carbohydrate structures. Still, it is unknown how this carbohydrate diversity affects the recognition by IgG antibodies produced as a response to viral infection.

### Results

We expressed the receptor binding domain (RBD) of the spike protein in three different mammalian cell lines and identified cell-line specific differences in the carbohydrate profile. Sialic acids were a common trait of RBD produced in Chinese hamster ovary cells, while fucose was more commonly found within the carbohydrate structures on RBD produced in Human embryonic kidney cells. Both sialic acids and fucose were found to modulate the IgG-reactivity against the RBD, with enhanced reactivity as a result following enzymatic removal of these specific sugar residues.

### Impact

The results highlight the importance of the carbohydrate profile in antibody reactivity against the RBD of SARS-CoV-2, and suggests the production cell line is of importance when producing recombinant proteins as vaccine candidates.

## Expanded view figure legends and tables

**Table EV1.**
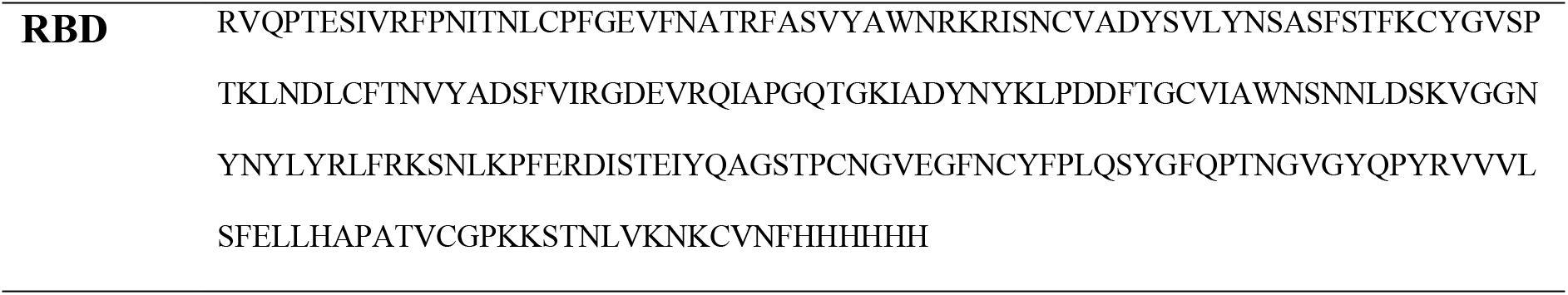
Amino acid sequences of the recombinant RBD including a C-terminal His-tag.

**Table EV2.** Percentage distribution between different amounts of antenna. The lower part of the table shows presence of LacDiNAc, degree of fucosylation and degree of sialylation on complex type glycans produced in CHO-S and HEK293F-cell lines.

|  | N331 |  | N343 |  |
| --- | --- | --- | --- | --- |
|  | CHO-S | HEK293F | CHO-S | HEK293F |
| mono-antennary | 4.3 | 4.8 | 7.8 | 2.3 |
| mono-antennary/bi-antennary<sup>†</sup> |  | 12.1 |  | 25.7 |
| bi-antennary | 52.2 | 52.5 | 68.5 | 60.4 |
| bi-antennary/tri-antennary<sup>†</sup> |  | 21.5 |  | 8.8 |
| tri-antennary | 26.3 | 5.5 | 19.5 | 1.9 |
| tri-antennary/tetra-antennary/repeats<sup>†</sup> | 17.1 | 3.6 | 4.3 | 0.8 |
| LacDiNAc* | 0.0 | 79.7 | 0.0 | 70.3 |
| Fucose* | 95.9 | 100.0 | 91.5 | 99.1 |
| Sialic acid* | 65.0 | 38.6 | 47.9 | 14.1 |

<sup>†</sup> Individual fragment ion spectra supporting the presence of either single HexNAc or DiHexNAc were observed for the glycoform of the same glycan composition, making it not always possible to assign an unambiguous
number of the antenna structures. For relative quantification purposes, those are presented as dual assignment structures.
\* The percentage is calculated in relation to total amount of complex structures.

**Table EV3.** Percentage of structures carrying zero to four fucose groups within the same structure showed for site N331 and N343 produced in CHO-S, HEK293F and Lec3.2.8.1-cells. The lower part of the table shows the fucosylation level within the observed glycoforms.

| # of Fucose residues | N331 |  |  | N343 |  |  |
| --- | --- | --- | --- | --- | --- | --- |
|  | CHO-S | HEK293F | Lec3.2.8.1 | CHO-S | HEK293F | Lec3.2.8.1 |
| 0 | 26.8 | 3.4 | 68.6 | 21.6 | 6.5 | 93.7 |
| 1 | 73.0 | 54.8 | 31.4 | 78.3 | 69.9 | 6.3 |
| 2 | 0.2 | 36.2 | 0.0 | 0.0 | 21.3 | 0.0 |
| 3 | 0.0 | 5.6 | 0.0 | 0.0 | 2.0 | 0.0 |
| 4 | 0.0 | 0.1 | 0.0 | 0.0 | 0.3 | 0.0 |
| Oligomannose* | 0.0 | 0.0 | 34.1 | 0.0 | 0.0 | 7.9 |
| Paucimannose* | 32.8 | 86.6 | 60.5 | 56.5 | 62.5 | 0.0 |
| Hybrid* | 94.3 | 100.0 | 0.0 | 55.2 | 60.4 | 0.0 |
| Complex* | 95.9 | 100.0 | 100.0 | 91.5 | 99.1 | 0.0 |

\* The percentage is calculated in relation of total amount of all observed glycoforms within the given glycan group, i.e. oligomannose, paucimannose, hybrid and complex. For the percentage distribution of the glycoforms at each site see Table 1.

**Table EV4.**
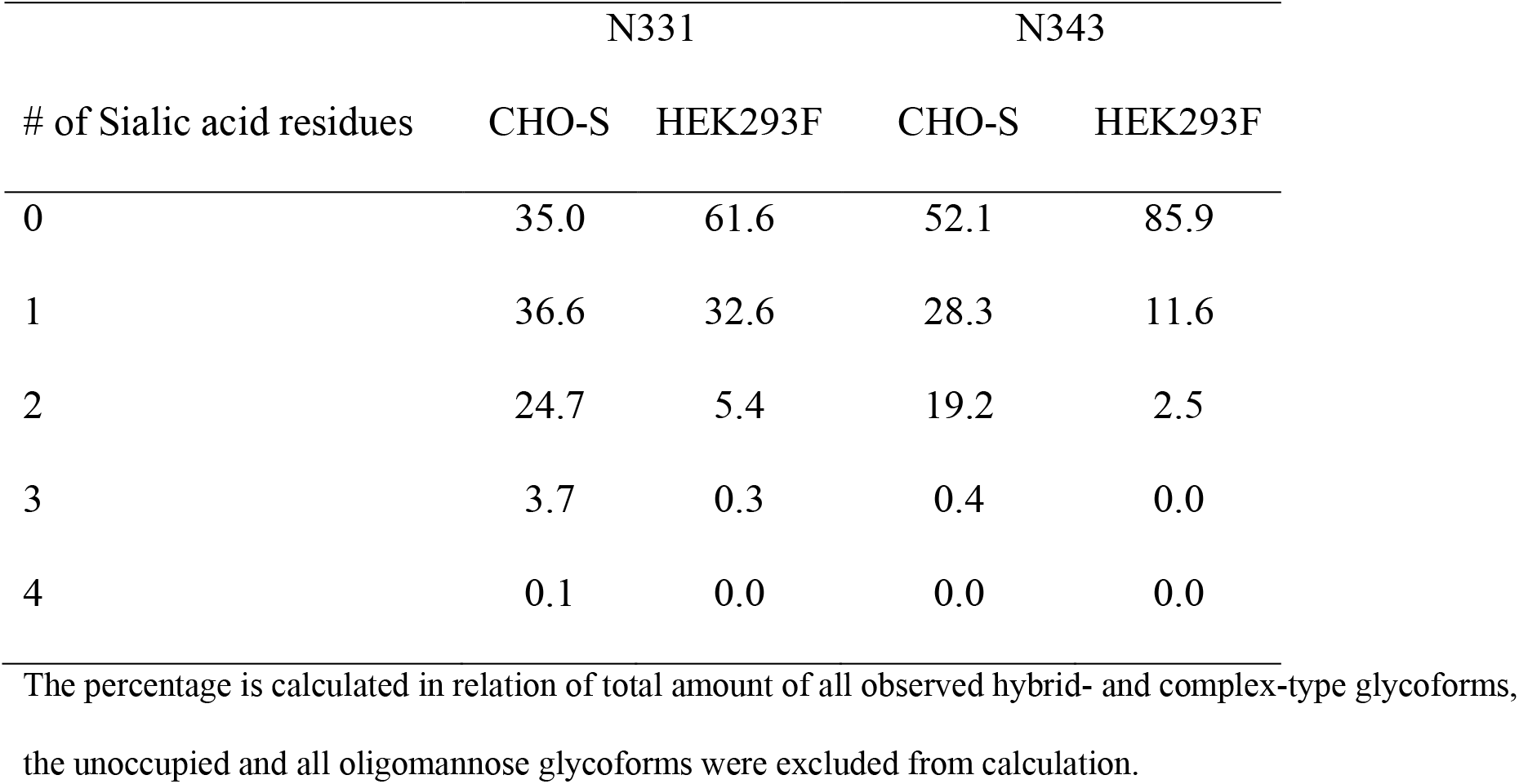
Percentage of detected structures carrying zero to four sialic acids within the same glycan structure.

**Table EV5.**
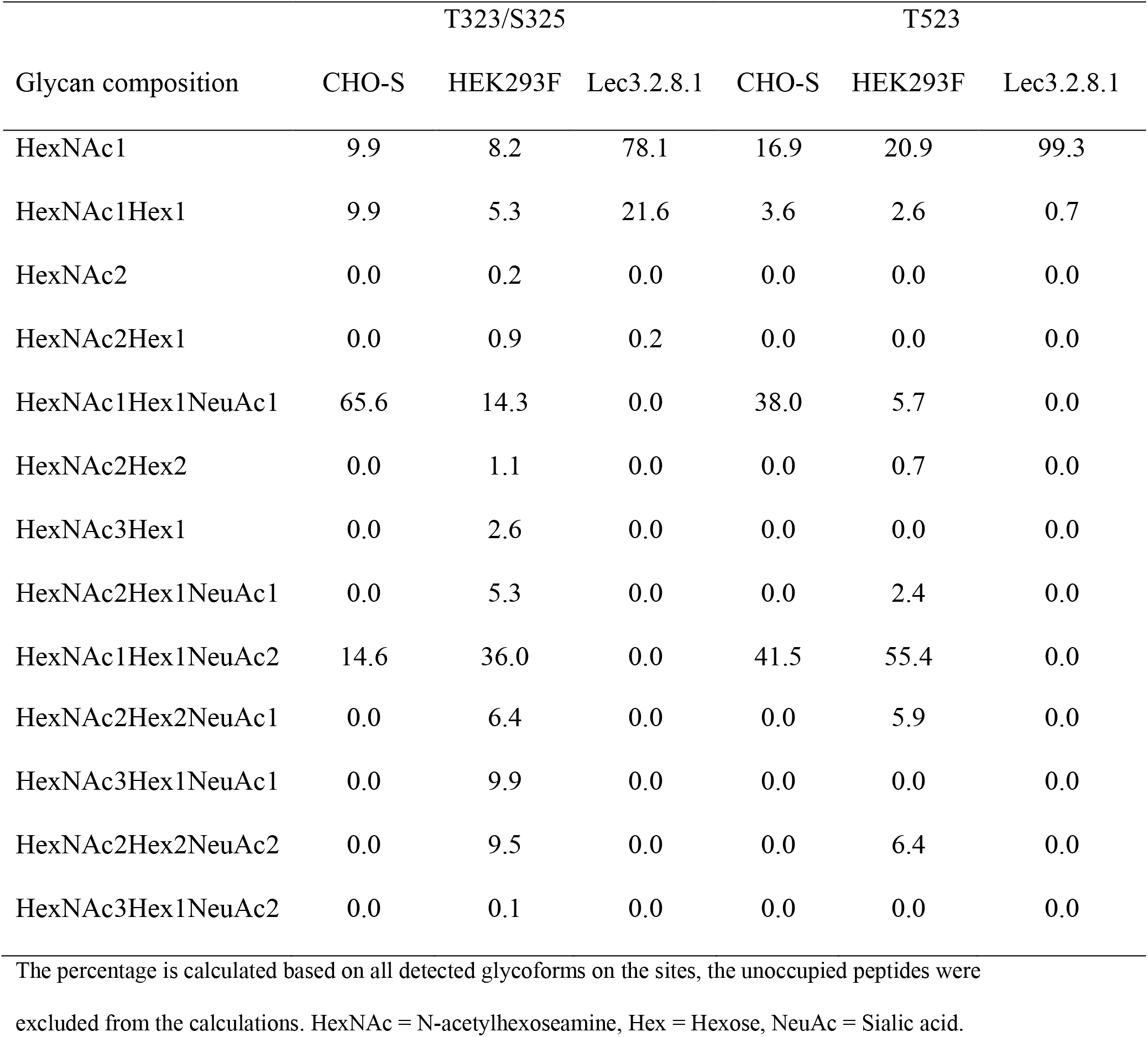
Percentage of O-linked structures at site T323/S325 and T523 when produced in CHO-S, HEK293F, and Lec3.2.8.1-cells.

**Table EV6.**
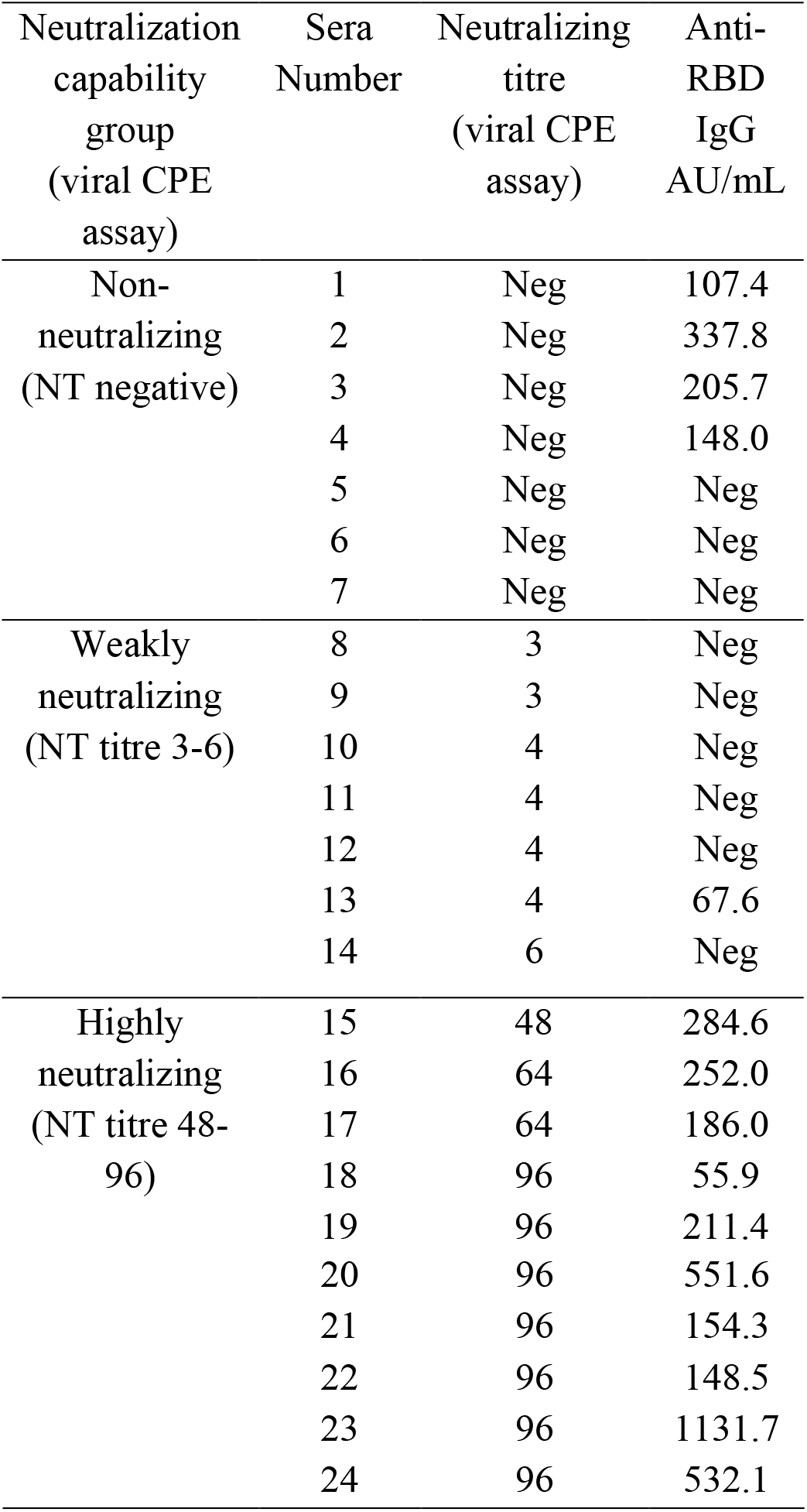
Characterization of convalescent sera. Neutralization capability determined using a viral CPE assay. Anti-RBD IgG levels as determined using an automated CMIA (values ≥ 50 AU/mL is considered positive). The neutralization capability group is stated as decided by the Viral CPE assay.

**Figure EV1.**
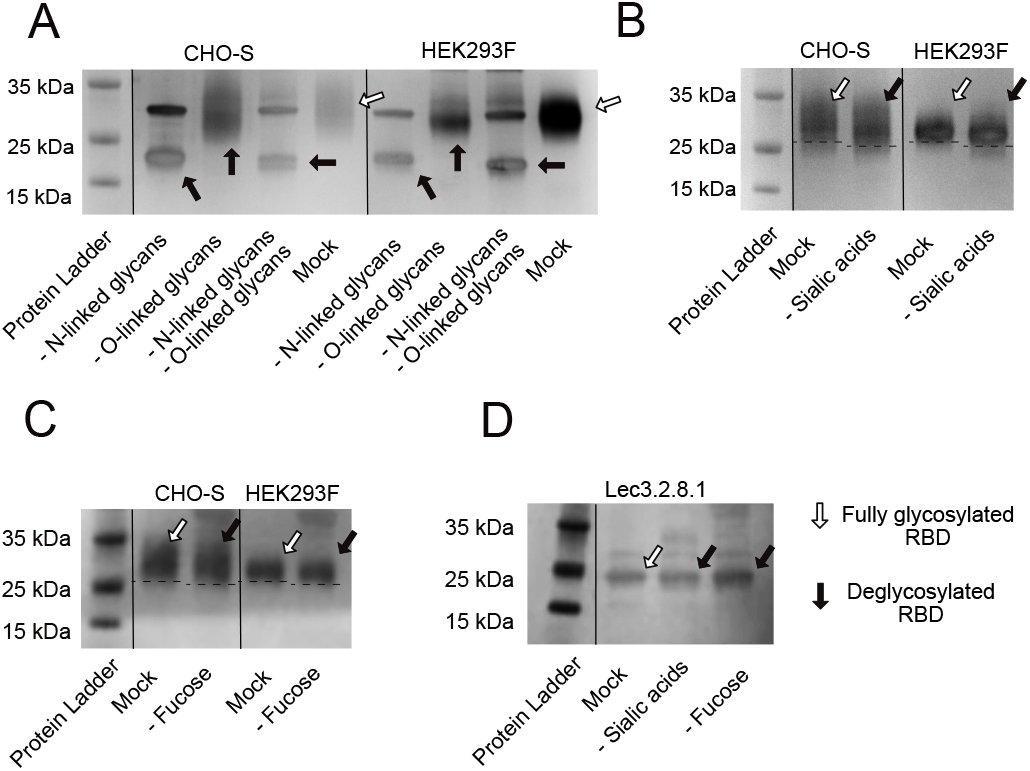
Silver stain of SDS page gel showing band size shift after deglycosylation. A. Band size shift after removal of N-linked, O-linked or both N-linked and O-linked glycans from RBD produced in CHO-S and HEK293F-cells. Open arrows represent fully glycosylated constructs while solid arrows represent the deglycosylated variants. Remaining bands originates from enzymes used in the deglycosylation-reaction. B. Band size shift following sialidase treatment of recombinant RBD produced in CHO-S and HEK293F-cells. C. Band size shift following fucosidase treatment of recombinant RBD produced in CHO-S and HEK293F-cells. D. Band size following sialidase and fucosidase treatment of recombinant RBD produced in Lec3.2.8.1-cells. Data information: The PageRuler™ Prestained Protein ladder is used as size comparison.

**Figure EV2.**
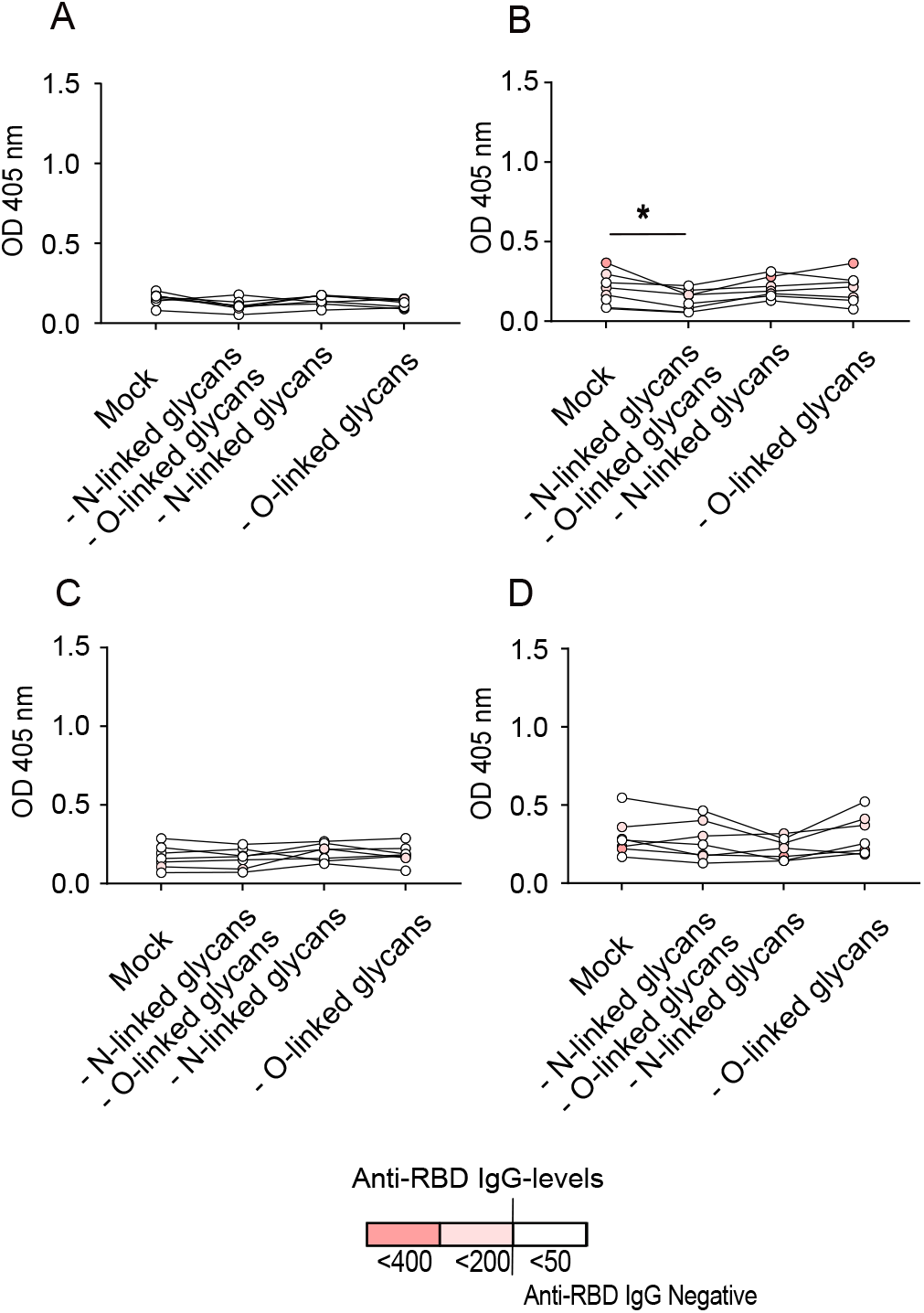
Serum reactivity against the fully glycosylated RBD (mock-treated) and against RBD following removal of N-linked, O-linked or both N-linked and O-linked glycans. A. Antibody reactivity of weakly neutralizing sera (NT nitre 3-6, n=7) against RBD produced in CHO-S-cells. B. Antibody reactivity of non-neutralizing sera (NT negative, n=7) against RBD produced in CHO-S-cells. C. Antibody reactivity of weakly neutralizing sera (NT nitre 3-6. n=7) against RBD produced in HEK293F-cells. D. Antibody reactivity of non-neutralizing sera (NT negative, n=7) against RBD produced in HEK293F-cells. Data information: Dark red colour symbolizes a serum with high levels of anti-RBD IgG, white colour indicates anti-RBD IgG-negative serum (<50 AU/mL). Statisitical analysis was performed with Wilcoxon matched-pair signed rank test. * = p < 0.05.

**Figure EV3.**
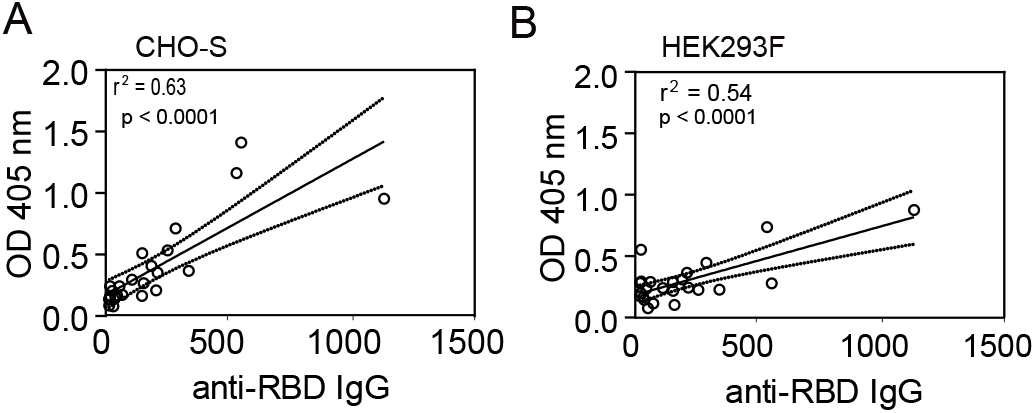
Correlation between anti-RBD IgG titres and OD (405 nm) as measured with ELISA using fully glycosylated RBD. A. RBD produced in CHO-S-cells. B. RBD produced in HEK293F cells. Data information: Pearson correlation coefficient analysis showed a strong correlation (p < 0.0001) between high anti-RBD IgG-levels and high OD-value for both CHO-S and HEK293F-produced constructs.

## References

Abbott RK, Crotty S (2020) Factors in B cell competition and immunodominance. Immunol Rev 296: 120–131

Allen JD, Chawla H, Samsudin F, Zuzic L, Shivgan AT, Watanabe Y, He W-t, Callaghan S, Song G, Yong P et al (2021) Site-Specific Steric Control of SARS-CoV-2 Spike Glycosylation. Biochemistry 60: 2153–2169

Antonopoulos A, Broome S, Sharov V, Ziegenfuss C, Easton RL, Panico M, Dell A, Morris HR, Haslam SM (2021) Site-specific characterization of SARS-CoV-2 spike glycoprotein receptor-binding domain. Glycobiology 31: 181–187

Bagdonaite I, Thompson AJ, Wang X, Søgaard M, Fougeroux C, Frank M, Diedrich JK, Yates JR, Salanti A, Vakhrushev SY et al (2021) Site-specific O-glycosylation analysis of SARS-CoV-2 spike protein produced in insect and human cells. bioRxiv

Brun J, Vasiljevic S, Gangadharan B, Hensen M, A VC, Hill ML, Kiappes JL, Dwek RA, Alonzi DS, Struwe WB et al (2021) Assessing Antigen Structural Integrity through Glycosylation Analysis of the SARS-CoV-2 Viral Spike. ACS Cent Sci 7: 586–593

Cavanagh D (1983) Coronavirus IBV: structural characterization of the spike protein. The Journal of general virology 64 (Pt 12): 2577–2583

Chen W, Stanley P (2003) Five Lec1 CHO cell mutants have distinct Mgat1 gene mutations that encode truncated N-acetylglucosaminyltransferase I. Glycobiology 13: 43–50

Chi X, Yan R, Zhang J, Zhang G, Zhang Y, Hao M, Zhang Z, Fan P, Dong Y, Yang Y et al (2020) A neutralizing human antibody binds to the N-terminal domain of the Spike protein of SARS-CoV-2. Science 369: 650–655

Croset A, Delafosse L, Gaudry J-P, Arod C, Glez L, Losberger C, Begue D, Krstanovic A, Robert F, Vilbois F et al (2012) Differences in the glycosylation of recombinant proteins expressed in HEK and CHO cells. Journal of Biotechnology 161: 336–348

Delmas B, Laude H (1990) Assembly of coronavirus spike protein into trimers and its role in epitope expression. Journal of virology 64: 5367–5375

Doering TL, Cummings RD, Aebi M (2015) Fungi. In: Essentials of Glycobiology, Varki A., Cummings R.D., Esko J.D., Stanley P., Hart G.W., Aebi M., Darvill A.G., Kinoshita T., Packer N.H., Prestegard J.H. et al (eds.) pp. 293–304. Cold Spring Harbor Laboratory Press

Copyright 2015-2017 by The Consortium of Glycobiology Editors, La Jolla, California. All rights reserved.: Cold Spring Harbor (NY)

Ghosh S, Dellibovi-Ragheb TA, Kerviel A, Pak E, Qiu Q, Fisher M, Takvorian PM, Bleck C, Hsu VW, Fehr AR et al (2020) β-Coronaviruses Use Lysosomes for Egress Instead of the Biosynthetic Secretory Pathway. Cell 183: 1520–1535.e1514

Grant OC, Montgomery D, Ito K, Woods RJ (2020) Analysis of the SARS-CoV-2 spike protein glycan shield reveals implications for immune recognition. Scientific reports 10: 14991

Gstöttner C, Zhang T, Resemann A, Ruben S, Pengelley S, Suckau D, Welsink T, Wuhrer M, Domínguez-Vega E (2021) Structural and Functional Characterization of SARS-CoV-2 RBD Domains Produced in Mammalian Cells. Anal Chem 93: 6839–6847

Ho JK-T, Jeevan-Raj B, Netter H-J (2020) Hepatitis B Virus (HBV) Subviral Particles as Protective Vaccines and Vaccine Platforms. Viruses 12: 126

Ju B, Zhang Q, Ge J, Wang R, Sun J, Ge X, Yu J, Shan S, Zhou B, Song S et al (2020) Human neutralizing antibodies elicited by SARS-CoV-2 infection. Nature 584: 115–119

Kellam P, Barclay W (2020) The dynamics of humoral immune responses following SARS-CoV-2 infection and the potential for reinfection. Journal of General Virology 101: 791–797

Li W, Moore MJ, Vasilieva N, Sui J, Wong SK, Berne MA, Somasundaran M, Sullivan JL, Luzuriaga K, Greenough TC et al (2003) Angiotensin-converting enzyme 2 is a functional receptor for the SARS coronavirus. Nature 426: 450–454

Li Y, Ma M-l, Lei Q, Wang F, Hong W, Lai D-y, Hou H, Xu Z-w, Zhang B, Chen H et al (2021) Linear epitope landscape of the SARS-CoV-2 Spike protein constructed from 1,051 COVID-19 patients. Cell Rep 34: 108915

Lou B, Li TD, Zheng SF, Su YY, Li ZY, Liu W, Yu F, Ge SX, Zou QD, Yuan Q et al (2020) Serology characteristics of SARS-CoV-2 infection after exposure and post-symptom onset. Eur Respir J 56

Nordén R, Halim A, Nyström K, Bennett EP, Mandel U, Olofsson S, Nilsson J, Larson G (2015) O-linked glycosylation of the mucin domain of the herpes simplex virus type 1-specific glycoprotein gC-1 is temporally regulated in a seed-and-spread manner. The Journal of biological chemistry 290: 5078–5091

Nordén R, Nilsson J, Samuelsson E, Risinger C, Sihlbom C, Blixt O, Larson G, Olofsson S, Bergstrom T (2019) Recombinant Glycoprotein E of Varicella Zoster Virus Contains Glycan-Peptide Motifs That Modulate B Cell Epitopes into Discrete Immunological Signatures. International journal of molecular sciences 20

Olofsson S, Blixt O, Bergstrom T, Frank M, Wandall HH, 2016. Viral O-GalNAc peptide epitopes: a novel potential target in viral envelope glycoproteins, Reviews in medical virology, 2015/11/03 ed., pp. 34–48.

Pinto D, Park YJ, Beltramello M, Walls AC, Tortorici MA, Bianchi S, Jaconi S, Culap K, Zatta F, De Marco A et al (2020) Cross-neutralization of SARS-CoV-2 by a human monoclonal SARS-CoV antibody. Nature 583: 290–295

Plotkin SA, Plotkin SA (2008) Correlates of Vaccine-Induced Immunity. Clinical Infectious Diseases 47: 401–409

Raska M, Takahashi K, Czernekova L, Zachova K, Hall S, Moldoveanu Z, Elliott MC, Wilson L, Brown R, Jancova D et al (2010) Glycosylation patterns of HIV-1 gp120 depend on the type of expressing cells and affect antibody recognition. The Journal of biological chemistry 285: 20860–20869

Reed LJ, Muench H (1938) A SIMPLE METHOD OF ESTIMATING FIFTY PER CENT ENDPOINTS12. American Journal of Epidemiology 27: 493–497

Rydyznski Moderbacher C, Ramirez SI, Dan JM, Grifoni A, Hastie KM, Weiskopf D, Belanger S, Abbott RK, Kim C, Choi J et al (2020) Antigen-Specific Adaptive Immunity to SARS-CoV-2 in Acute COVID-19 and Associations with Age and Disease Severity. Cell 183: 996–1012.e1019

Sanda M, Morrison L, Goldman R (2021) N- and O-Glycosylation of the SARS-CoV-2 Spike Protein. Analytical Chemistry 93: 2003–2009

Shajahan A, Supekar NT, Gleinich AS, Azadi P (2020) Deducing the N- and O-glycosylation profile of the spike protein of novel coronavirus SARS-CoV-2. Glycobiology

Shental-Bechor D, Levy Y (2008) Effect of glycosylation on protein folding: a close look at thermodynamic stabilization. Proceedings of the National Academy of Sciences of the United States of America 105: 8256–8261

Tortorici MA, Beltramello M, Lempp FA, Pinto D, Dang HV, Rosen LE, McCallum M, Bowen J, Minola A, Jaconi S et al (2020) Ultrapotent human antibodies protect against SARS-CoV-2 challenge via multiple mechanisms. Science 370: 950–957

Wang C, Li W, Drabek D, Okba NMA, van Haperen R, Osterhaus A, van Kuppeveld FJM, Haagmans BL, Grosveld F, Bosch BJ (2020) A human monoclonal antibody blocking SARS-CoV-2 infection. Nature communications 11: 2251

Watanabe Y, Allen JD, Wrapp D, McLellan JS, Crispin M (2020a) Site-specific glycan analysis of the SARS-CoV-2 spike. Science 369: 330–333

Watanabe Y, Allen Joel D, Wrapp D, McLellan Jason S, Crispin M (2020b) Site-specific glycan analysis of the SARS-CoV-2 spike. Science 369: 330–333

Yang J, Wang W, Chen Z, Lu S, Yang F, Bi Z, Bao L, Mo F, Li X, Huang Y et al (2020) A vaccine targeting the RBD of the S protein of SARS-CoV-2 induces protective immunity. Nature

Zhao J, Yuan Q, Wang H, Liu W, Liao X, Su Y, Wang X, Yuan J, Li T, Li J et al (2020) Antibody Responses to SARS-CoV-2 in Patients With Novel Coronavirus Disease 2019. Clinical infectious diseases: an official publication of the Infectious Diseases Society of America 71: 2027–2034

